# Platelet and releasate lipidomics identify novel platelet activating mechanisms and ether-linked lysophosphatidylcholine and phosphatidylethanolamine 38:7 as predictors for platelet reactivity

**DOI:** 10.1101/2025.10.09.681267

**Authors:** Kang-Yu Peng

**Affiliations:** Sydney Mass Spectrometry, Charles Perkins Centre, the University of Sydney, Camperdown, New South Wales, Australia; The Heart Research Institute, Newtown, New South Wales, Australia

## Abstract

The activation of platelets and consequent thrombi formation are hallmark events for cardiovascular diseases. Both the secretion of lipid (in the platelet releasate) and intra-platelet lipid signaling have been shown to be important modulators to platelet activity and thrombosis. However, the relationship between platelet and releasate lipid profiles and how they respond to key agonists remain less clear.

With a purposely designed human platelet targeted lipidomics platform which detected more than 550 platelet and releasate lipid species encompassing >30 lipid classes/subclasses, I presented in this study major changes to platelet lipidome triggered by acute and prolonged thrombin and collagen stimulations. I have demonstrated that overall lipid release from platelets was suppressed by collagen at the acute phase, a phenomenon that to my knowledge has yet to be reported. Additionally, prolonged thrombin treatment did not cause further production of lipid mediators. It nevertheless triggered an overall release of lipid into the surrounding (as releasate), including several proinflammatory and apoptotic lipids e.g., DG, ceramide, S1P, PA.

Correlation tests between platelet/releasate lipid profiles and platelet surface markers P-selectin and PAC1 suggested that several di-/polyunsaturated phospholipid species and CoQ negatively correlated with collagen-stimulated PAC1 expression and PI 40:4(b) positively correlated with collagen-stimulated P-selectin expression. Finally, our study identified one lipid class, LPC-O, and one lipid species, PE 38:7, as potential predictors for the reactivity of platelet, with platelet LPC-O level positively correlated with collagen-stimulated increment of surface PAC1 expression and PE 38:7 correlated negatively with the increase in surface P-selectin in response to thrombin treatment.

## Introduction

Platelets (thrombocytes) are tiny and anucleated cells which play important roles in wound healing process (haemostasis) but are also considered a main culprit for fatal and debilitating cardiovascular diseases (CVD) such as occlusive stroke, pulmonary embolism and coronary artery disease, with pathogenesis often involving thrombus formation triggered by hyperactivation of platelets^1,2^.

Lipid has been shown to be a key modulator for platelet activity^1,2^. Recent advances in LC/MS-based lipidomics, i.e., systemic analysis of the “lipidome”^3,4^, provide novel and wholistic insights into the role of lipid in platelet functions. Importantly, ex vivo administration of platelet activating agents such as thrombin and collagen has been widely used to simulate how lipidome responded to various stimulations and how this impacted platelet functions^5–8^.

With all the exciting discoveries, earlier platelet lipidomics studies nonetheless were often limited by one or more of the following factors (1) small sample sizes, typically 3 to 5 replicates only; (2) lack of comprehensive releasate lipid profiles alongside platelet lipid profiles, (3) absence of studies regarding the effect of prolonged stimulation on platelet and releasate lipidomes, a likely scenario under the disease setting but rarely studied ex vivo, and (4) missing links between platelet and releasate lipidomes and platelet reactivity to stimulations^5–8^.

To address these shortcomings, A dedicated targeted (UHPLC-MS/MS) lipidomics platform was developed and utilized to analyse human platelet and releasate samples. More than 550 lipid species in collected platelet and releasate samples from 15 healthy donors have been reported here, making this work one the most comprehensive platelet targeted lipidomics studies to date. Profound changes to platelet and releasate lipidomes in response to stimulations have been identified. Concomitant mapping of the increase, decrease, secretion and retention of platelet and releasate lipid species in reaction to acute (10 minutes) and prolonged (60 minutes) thrombin and collagen stimulations was also carried out. Finally, I studied the relationship between platelet and releasate lipidomes and two surface markers of platelet activation, PAC1 and P-selectin, and identified correlations between several lipid species and classes and platelet surface expression of these markers.

## Abbreviations

8(9)-EET: 8,9-epoxyeicosatrienoic acid; 12-HETE 12-hydroxyeicosatetraenoic acid; AA: arachidonic acid; CAR: acylcarnitine; CE: cholesteryl ester; Cer: ceramide; Chol: cholesterol; CL: cardiolipin; CoQ: coenzyme Q; DG: diacylglycerol; dhCer: dihydroceramide; GM3: GM3 ganglioside; Hex2Cer: dihexosylceramide; HexCer: monohexosylceramide; LC/MS: liquid chromatography tandem mass spectrometry; LM: lipid mediators; LPC: lysphosphatidylcholine; LPC-O: alkyl lysophosphatidylcholine; LPC-P: alkenyl lysophosphatidylcholine; LPE: lysophosphatidylethanolamine; LPE-P: alkenyl lysophosphatidylethanolamine; LPI: lysophosphatidylinositol; LPS: lysophosphatidylserine; PA: phosphatidic acid; PAF: platelet activating factor; PC: phosphatidylcholine; PC1: principal component 1; PC2: principal component 2; PCA: principal component analysis; PC-O: alkyl phosphatidylcholine; PC-P: alkenyl phosphatidylcholine; PE: phosphatidylethanolamine; PE-O: alkyl phosphatidylethanolamine; PE-P: alkenyl phosphatidylethanolamine; PG: phosphatidylglycerol; PI: phosphatidylinositol; PL: phospholipid; PLA2: phospholipase A2; PS: phosphatidylserine; S1P: sphingosine-1-phosphate; SM: sphingomyelin; Sph: sphingosine; SRM: single reaction monitoring; TG: triacylglycerol; TXB2: thromboxane B2.

## Results

### Overview of healthy human platelet and releasate lipid profiles

Washed platelets isolated from 15 healthy donors of diverse age, gender and ethnicity backgrounds by sequential centrifugations were used in this study. To study the acute and chronic effects of stimuli on platelet and releasate lipidomes, the washed platelets were treated with collagen (10 μg/mL), low (0.2 U/mL) or high dose (1.0 U/mL) thrombin for 10 or 60 minutes. Furthermore, the levels of platelet surface PAC1 and P-selectin (CD62p) were determined by flow cytometry^9^. Platelet and releasate lipid extractions were subsequently performed using the butanol and methanol (BUME) approach^10^ (Figure 1).

**Figure 1.**
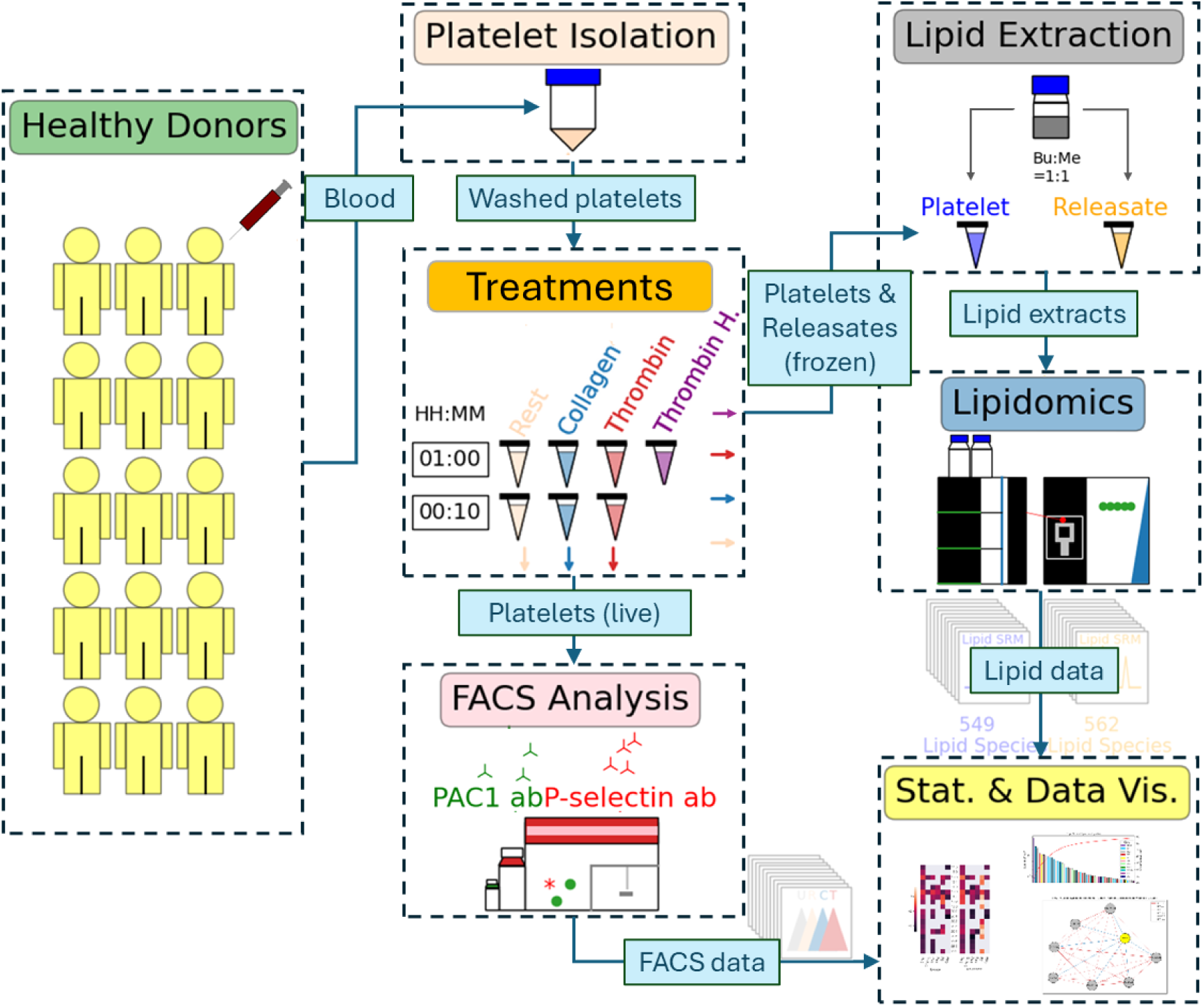
Overview of study design. Blood from 15 healthy adult donors was collected for platelet isolation, as described in the “Materials and Methods” section. Platelets were then treated without (Rest) or with 10 μg/mL collagen (Collagen), 0.2 U/mL thrombin (Thrombin) or 1.0 U/mL thrombin (Thrombin H.) for 10 or 60 minutes before being palleted by centrifugation. The pellet (platelets) and supernatant (releasates) were respectively collected and immediately stored under −80°C. FACS analysis for platelet surface markers was also performed using live platelets staining with FITC-conjugated PAC1 or APC-conjugated P-selectin antibodies followed by 10-minute collagen (10 μg/mL) or thrombin (0.2 U/mL) stimulation. Extracted platelet and releasate samples were subsequently analysed with the targeted lipidomics platform. Overall, 549 and 562 lipid species were detected in platelet and releasate samples, respectively, with all SRM chromatograms underwent careful manual curations. Relevant statistical analysis and data visualization were performed on the lipid and FACS data.

Our targeted lipidomics platform detected 549 and 562 lipid species in human platelet and releasate, respectively. As shown in Figure 2A, diverse lipid classes encompassing phospholipids, lysophospholipids, sphingolipids, lysosphingolipids, cholesterol, neutral lipids and others (lipid mediators, coenzyme Q and acylcarnitine) were observed in resting platelet lipidome. Notably, the concentrations of these lipid classes varied by ~10^4^ folds between the most and least abundant lipid classes found in platelets. Lipid classes that existed in platelet were similarly found in the releasate. However, the low abundance lipid classes were scarcer in the releasate than platelet, making the maximal between-class concentration difference nearly 10^5^ folds (Figure 2B).

**Figure 2.**
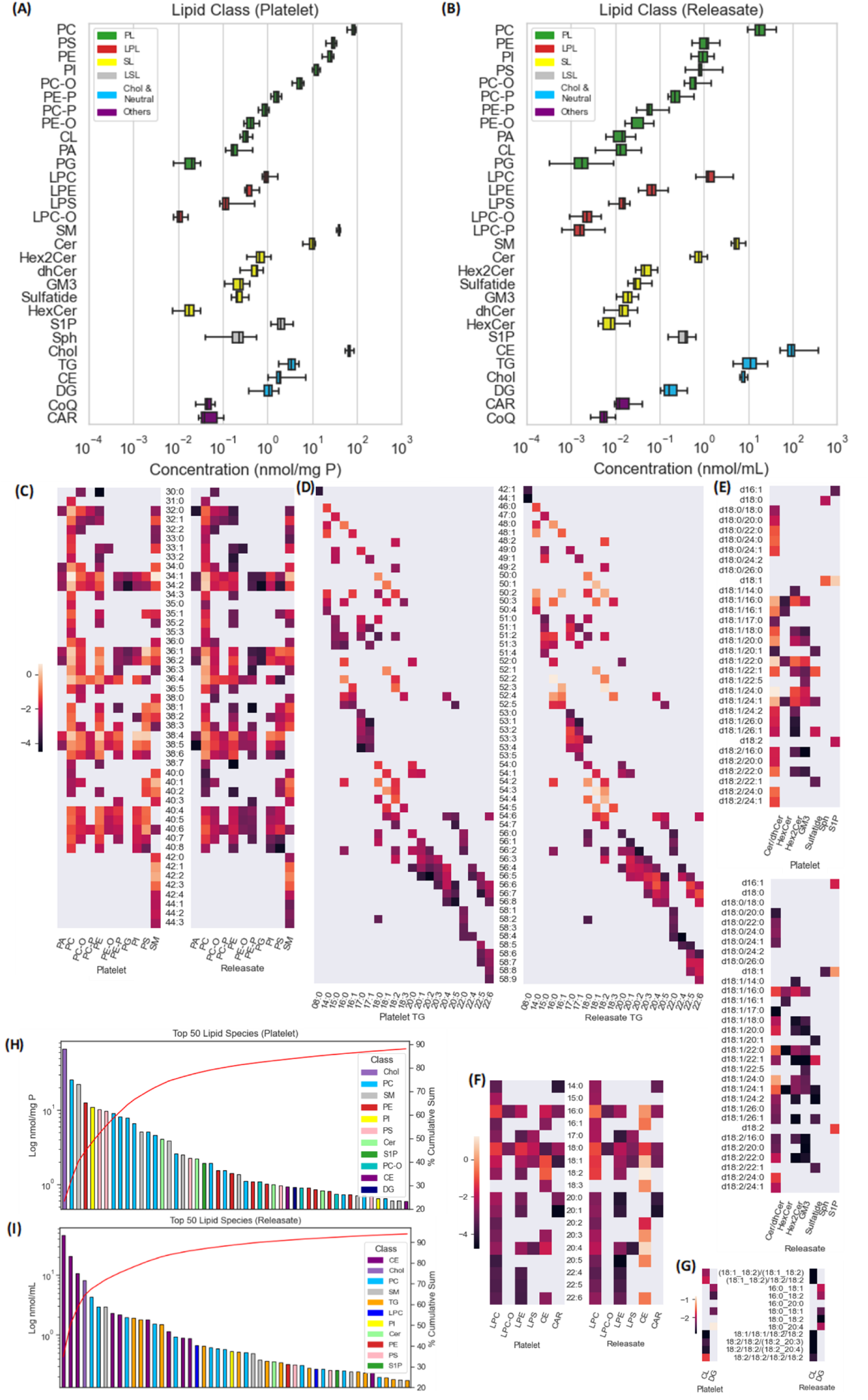
Human platelet and releasate lipidomics. Resting platelet (A) and releasate (B) lipid class profiles were shown as box plots (abundances high to low within lipid categories). Heatmap representations for the phospholipid and SM (C), TG (D), sphingo- and lysosphingolipid (E), lysophospholipid, CE and CAR (F) and CL and DG (G) detected in platelet and releasate were likewise shown, with logged concentrations indicated by colour bars. Bar plots for top 50 most abundant lipid species in platelet (H) and releasate (I), with red curves indicating % cumulative sums of total lipid (lipid mediators excluded).

Resting platelet and releasate lipidomes were further presented based on lipid classes and/or the fatty acyl chains and saturation levels in Figure 2C-G. Heatmap representations demonstrated that the difference between platelet and releasate lipidomes occurred mainly at the lipid class rather than the lipid species level, i.e., lipid species in a class could be consistently lower (e.g., PE, PE-O, PE-P, Cer/dhCer, Hex2Cer, GM3, CL) or higher (e.g., CE, TG) in the releasate than in the lysate. Nonetheless, within class composition was largely maintained. For platelets, the most abundant lipid species were cholesterol, PC, SM, PE, PI, PS and Cer (Figure 2H). Whereas the more abundant releasate lipid species were the CE, cholesterol, PC, SM, TG and LPC species (Figure 2I). The top 50 lipid species covered almost 90% of the platelet lipid. In the releasate, this number was roughly 95% (Figure 2H&I).

### Effects of acute and prolonged stimulations on platelet and releasate lipidomes

Principal component analysis (PCA) provided an overview of the effects of acute and prolonged collagen and thrombin treatments on platelet and releasate lipidomes (Figure 3A-D). Both collagen- and thrombin-activated platelets displayed differences in their lipid profiles compared with resting platelets, with thrombin exerting a stronger effect than collagen. Prolonged incubation for 60 minutes did not induce substantial differences vs 10-minute incubation in platelet lipids, for both collagen and thrombin treatments. High thrombin dose moderately increased the shift at 60-minute time point when compared to the low thrombin dose (Figure 3 A&B).

**Figure 3.**
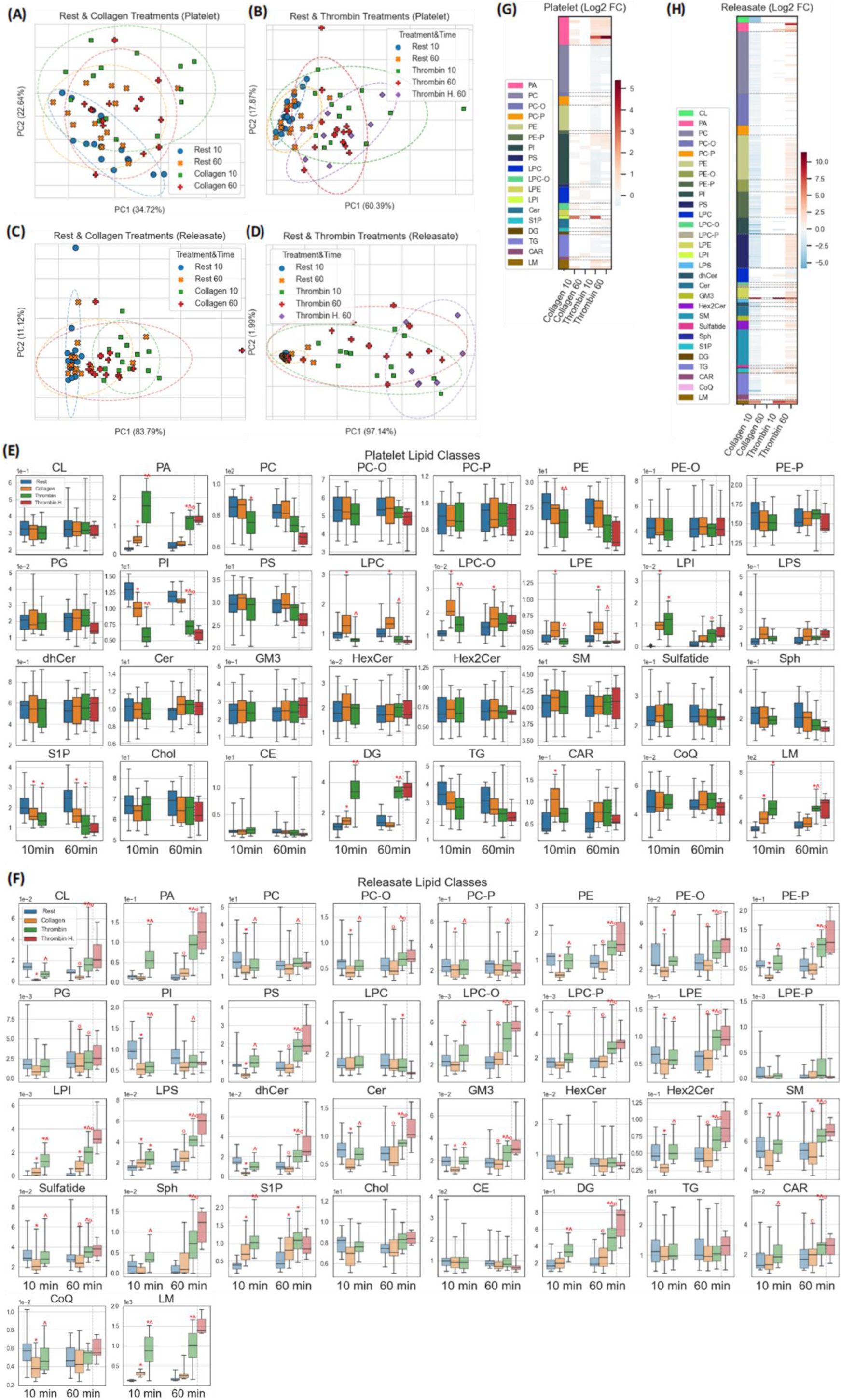
Collagen- and thrombin-induced alterations to platelet and releasate lipid profiles. Isolated platelets were treated without (Rest) or with 10 μg/mL collagen (Collagen) or 0.2 U/mL (Thrombin) or 1.0 U/mL thrombin (Thrombin H.) for 10 or 60 minutes, followed by lipidomics analysis (n=14). Overviews of acute and prolonged effects of thrombin and collagen on platelets and releasate were achieved by principal component analysis (PCA) (A-D). Effects of thrombin and collagen on platelet and releasate lipid classes were shown as boxplots (E&F). Significantly altered platelet (G) and releasate (H) lipid species were represented by heatmaps with the colour bars indicating logged median fold changes. *: p<0.05 vs time matched resting samples; ^: p<0.05 vs time matched collagen treatment; ᵒ: p<0.05 between two time points; Except for PCA, thrombin H. group was excluded when performing statistical analysis due to unmatched sample size.

Collagen and thrombin nonetheless triggered more pronounced changes to releasate lipid profiles. As demonstrated in Figure 3C, 10-minute collagen treatment induced a clear deviation from the time-matched resting samples, mainly along PC1. Prolonged incubation somewhat reversed this effect. Acute thrombin induced an even stronger effect than collagen and prolonged treatment, unlike collagen, did not result in a reversal. When compared with acute thrombin treatment, further deviation caused by prolonged treatment more evidently occurred along PC2 axis, suggesting involvements of a different set of features from the acute treatment. A high thrombin dose likewise induced a stronger shift toward right along PC1 (Figure 3D).

Detailed effects of collagen and thrombin on platelet lipid classes were presented as boxplots and shown in Figure 3E. Among phospholipids, the strongest effect occurred to PA (increase) and PI (decrease), with thrombin generally exerting more prominent changes than collagen. The accumulation of several lysophospholipid classes (LPC, LPC-O, LPE) were much more prominent in the collagen-stimulated groups than the thrombin-stimulated groups, with LPI being an exception. The only platelet sphingolipid class that changed significantly in response to the stimulations was S1P, which decreased with both collagen and thrombin treatments. Both collagen and thrombin treatments caused increments to platelet DG, but this effect was more pronounced with thrombin. Platelet CAR increased significantly with 10-minute collagen treatment. Finally, LM increased significantly with both collagen and thrombin treatments. However, the effect of thrombin was stronger and appeared to last longer (Figure 3E).

Likewise, I examined how collagen and thrombin altered the releasate lipid profile. Overall, the effects of both collagen and thrombin on the releasate were more pronounced than those on the platelet. Only 5 lipid classes were not impacted significantly by any treatments at all (LPE-P, HexCer, Chol, CE and TG). Acute collagen treatment surprisingly lowered many lipid classes, encompassing phospholipid (CL, PC, PC-O, PC-P, PE, PE-O, PE-P, PI, PS), lysophospholid (LPE), sphingolipid (dhCer, Cer, GM3, Hex2Cer, SM, Sulfatide) and CoQ, which were partially reversed after prolonged treatment. Only LPI, S1P and LM increased significantly in response to both acute and prolonged collagen treatments. Except for PI, thrombin treatment generally increased releasate lipid. Notably, many more releasate lipid classes increased significantly in reaction to prolonged thrombin treatment (CL, PA, PC-O, PE, PE-O, PE-P, PS, LPC-O, LPC-P, LPE, LPI, LPS, dhCer, Cer, GM3, Hex2Cer, SM, Sph, DG and CAR) than acute treatment (Figure 3F).

At platelet lipid species level, thrombin exerted much more pronounced changes than collagen, with PA 38:1 (10 & 60 min, thrombin) and LPI 18:0 (10 min) increasing most dramatically in terms of fold changes (Figure 3G). 10-minute Collagen treatment suppressed the release of a large number of lipid species, with LPI 18:0, S1Ps and the lipid mediators being notable exceptions. For thrombin treatments, releasate lipid was mainly increased, especially for various PA species, LPI 18:0, DG 18:0_20:4 and the lipid mediators. Numerous lipid species were released following prolonged thrombin treatment (Figure 3H).

### Mapping stimulated platelet and releasate lipidomes

The closed nature of the ex vivo model used in this study made it possible to study the secretion and retention of platelet lipids and at the same time identify lipids potentially being metabolized in reaction to stimulations. To interrogate this, I first converted pmol/mg protein (platelet) and pmol/mL (releasate) data to total amount of lipid generated by 8×10^7^ platelets and in 200 μL releasate (new unit: 0.01 femtomole). Log10 median differences between treatments and rest (Δ) at the same time points (absolute values) were then plotted as Figure 4A-D, with 4 quadrants in each plot indicating changes in the two types of samples. From these plots, it was evident that platelet and releasate lipidomes, regardless of treatments, were closely related to each other, as the distribution of dot points, indicative of lipid species, generally follows the two diagonal lines. Two relationships exist here: (1) Δlipid in the releasate was positively correlated with the increase (1^st^ quadrant) or decrease (3^rd^ quadrant) of platelet lipid, or (2) Δlipid in the releasate was negatively correlated with the increase (4^th^ quadrant) or decrease (2^nd^ quadrant) of platelet lipid (Figure 4A-D & Supplementary Table S4).

**Figure 4.**
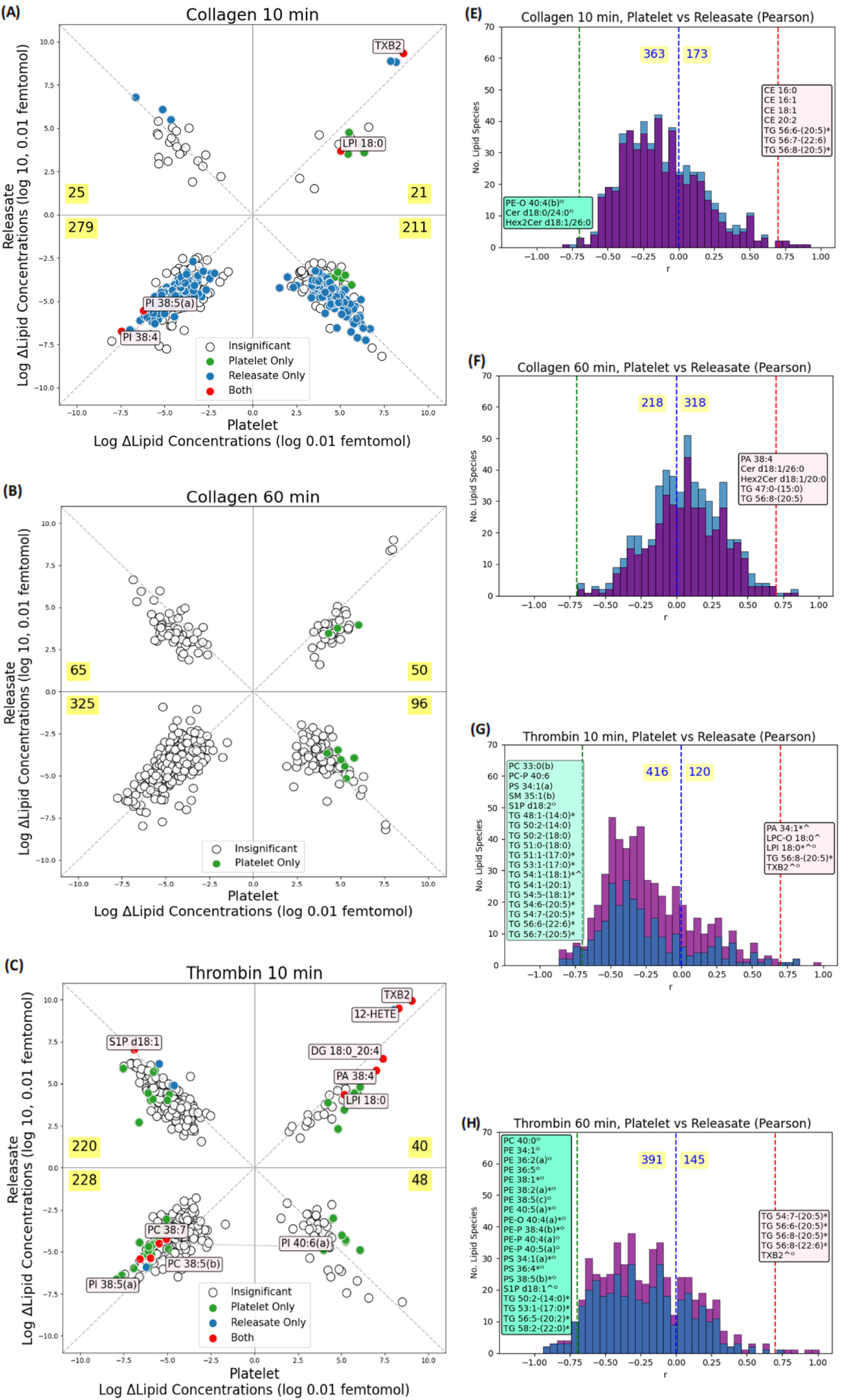
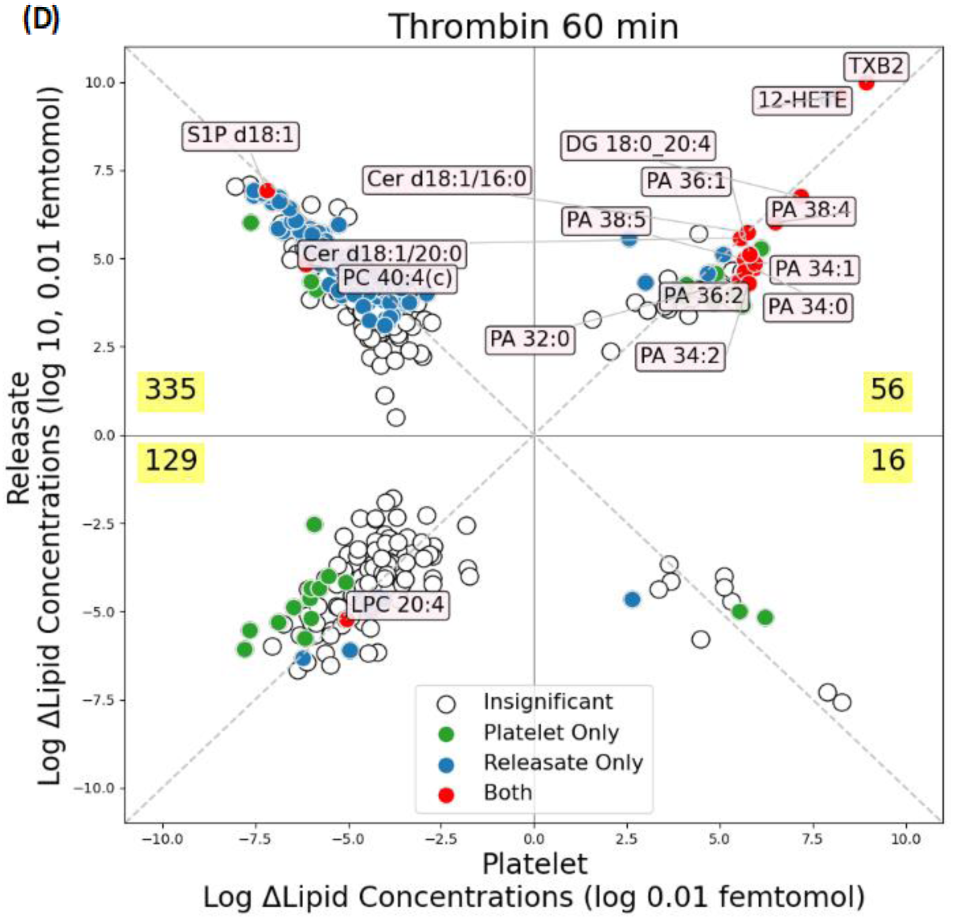
Delineating the relationship between platelet and releasate lipidomes in response to collagen and thrombin stimulations. Data were presented as total in vitro platelet and releasate lipid for individual lipid species. Log10 median values for the differences^#^ in lipid species between Collagen and Rest groups (log Δlipid concentrations) at 10 and 60 minutes (A&B) and those between Thrombin and Rest groups at 10 and 60 minutes (C&D) have been plotted, with significant lipid species (vs Rest) labelled. The highlighted values indicated the numbers of lipid species falling in specific quadrants. Pearson correlation tests between Δplatelet and Δreleasate at lipid species level were summarised as histogram plots (3E-H). Red and green lines indicated r>0.7 & r<−0.7, respectively, and only lipid species above the thresholds were listed. Blue lines (r=0) separate positive and negative correlations between the platelet and releasate data, with the numbers of positively and negatively correlated lipid species shown on both sides. For the histogram part, the purple bars indicate negative differences values in releasate (i.e., lipid species that were lower with the treatment in releasate), whereas the blue bars indicate positive differences values in releasate. #: greater or less than 0 for both the platelet and releasate axes indicated the increase/decrease post collagen or thrombin treatment in Figure 4A-4D. For instance, lipid species in the second quadrant (−,+) were decreased in platelet but increased in releasate in response to specific treatment. Absolute values for difference were used before log. Furthermore, the unit applied here was “0.01 femtomole” to ensure all the values, prior to log, were >1 so that taking log did not result in negative values. *: p<0.05, Benjamini-Hochberg corrected Pearson correlation tests; ^: p<0.05, treatment vs rest in platelet; °: treatment vs rest in releasate.

Acute collagen treatment appeared to lower the secretion of most lipid species detected (490). Only TXB2 and LPI 18:0 were significantly increased in both platelet and releasate. 2 polyunsaturated PI species, PI 38:4 and PI 38:5(a), were significantly decreased in both platelet and releasate (Figure 4A). This acute effect of collagen was to some extents reversed with prolonged treatment. Less lipid species decreased in the releasate (421) and hardly any significant differences vs the resting group were observed at 60-minute (Figure 4B). Furthermore, lipid species localized in 3^rd^ and 4^th^ quadrants, when compared to acute treatment, appeared to be elevated within their respective quadrants. A phenomenon that again indicates that overall secretion at 60-minute was not suppressed as much as 10-minute timepoint (Figure 4A&B).

Acute thrombin treatment, by contrast, presented a distinct lipid profile from collagen treatment. The fact that most lipid species fell within the 2^nd^ and 3^rd^ quadrants suggested that the predominant effects of thrombin was to enhance the secretion of platelet lipid (2^nd^ quadrant) and to lower some platelet lipid and therefore caused concomitant low releasate lipid levels (3^rd^ quadrant). On top of increased TXB2 and decreased polyunsaturated PI species similar to acute collagen treatment, acute thrombin treatment increased 12-HETE, DG 18:0_20:4 and PA 38:4 in both platelet and releasate. It also significantly decreased platelet and releasate polyunsaturated PC (PC 38:5(b) and PC 38:7). In addition, S1P d18:1 was significantly lower in platelet but higher in releasate and fell closely to the diagonal line, suggesting that this lipid species was released from platelet with thrombin stimulation (Figure 4C). 60-minute prolonged thrombin treatment furthered the release of many lipid species, as the numbers of lipid species in both 1^st^ and 2^nd^ quadrants increased dramatically. Many PA species were elevated in both platelet and releasate. Notably, two platelet and releasate ceramide species (Cer d18:1/16:0 and Cer d18:1/20:0) were also increased with prolonged thrombin treatment. A huge number (335) of lipid species increased significantly in releasate with simultaneous drops in their platelet contents. Among them, S1P d18:1 and PC 40:4 (c) were significantly different in both platelet and releasate. Finally, LPC 20:4 decreased significantly in both platelet and releasate in response to prolonged thrombin treatment (Figure 4D).

Pearson correlation tests were subsequently performed between platelet and releasate lipid species using the difference data (Δ). Relatively few (≤ 25 regardless of treatment) platelet and releasate lipid species showed highly positive (r > 0.7) or negative (r < −0.7) correlation (Figure 4E-H). Some of these lipid species belonged to lipid classes which were mostly insignificantly different between treatments and the rest samples (e.g., TG, CE). More platelet and releasate lipid species were highly correlated with thrombin treatment than collagen. With acute thrombin treatment, platelet and releasate PA 34:1 and LPC-O 18:0, two lipid species that significantly increased in platelet, were also highly and positively correlated between the two sample types. Platelet and releasate LPI 18:0 and TXB2, which were significantly elevated in both sample types, were also highly and positively correlated. S1P d18:2 (significantly increased in releasate) and TG 54:1-(18:1) (significantly increased in platelet) showed high negative correlations (Figure 4G). With prolonged thrombin treatment, the most noteworthy difference was that several phospholipid species that were only significantly elevated in releasate, especially PE, PE-O, PE-P and PS, became highly and negatively correlated between platelet and releasate, indicating the release of these lipid species (Figure 4H).

### Relationship between platelet surface markers and platelet and releasate lipids

Surface expressions of P-selectin and PAC1 are hallmark events for platelet activation. As demonstrated by FACS analysis, the level of P-selectin expression was significantly elevated by acute thrombin treatment (Figure 5A). Significant PAC1 expression could be seen with both collagen and thrombin stimulations (Figure 5B). Surface expressions of these two widely recognized markers for platelet activation nevertheless correlated poorly with each other regardless of the treatments (Figure 5C&D).

**Figure 5.**
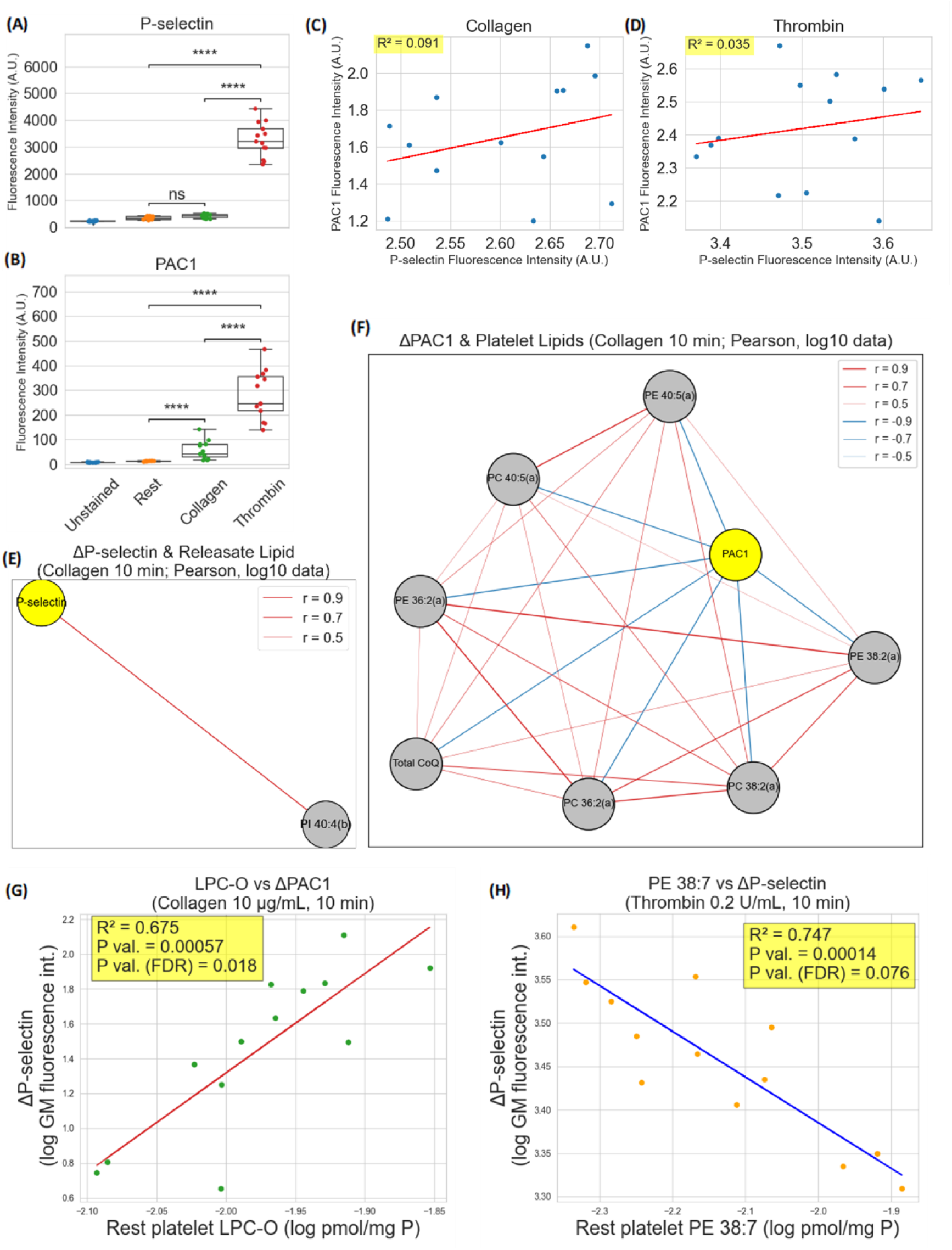
Correlations between platelet and releasate lipid profiles and platelet surface markers. The levels of P-selectin and PAC1 stains in unstained, rest, collagen and thrombin treated samples were presented in Figure 5A and 5B, with significances between the groups determined by Bonferroni corrected Mann-Whitney-Wilcoxon test. Network representations demonstrate the relationship between lipid species/classes that are significantly correlated (p < 0.05) or having strong tendencies toward significance (0.05 ≤ p < 0.06) with differences (Δ) to P-selectin and PAC1 levels (vs rest) in reaction to collagen stimulation for 10 minutes, as determined by Pearson correlation tests (logged data) (E&F). The respective levels of LPC-O and PE 38:7 in resting platelets were significantly correlated with PAC1 expression upon collagen stimulation and P-selectin expression upon thrombin stimulation, as determined by linear regression using logged data (G&H). ****: p ≤ 10^−4^, Mann-Whitney-Wilcoxon test with Bonferroni correction

To elucidate the relationship between platelet and releasate lipid and the expression of these markers for platelet activation upon stimulations, Pearson correlation tests were performed on each platelet/releasate lipid species and class against net increase in P-selectin or PAC1 levels, indicated by fluorescence intensities, post collagen or thrombin treatments (logged data; 10-minute treatment). Out of the >500 lipid species and >30 lipid classes in platelet and releasate, very few lipid species and only one lipid class were significantly or nearly significantly correlated with these platelet activation markers. Releasate PI 40:4(b) was the only lipid species highly and positively correlated (r = 0.89) with net increase in P-selectin surface presentation induced by collagen (Figure 5E). One platelet lipid class, CoQ, was negatively correlated with the increment to PAC1 (r = −0.81). Moreover, a number of di- and polyunsaturated (36:2, 38:2 and 40:5) PC and PE species in platelet also showed strong tendency toward significantly negative correlations (all with r < −0.81) with net PAC1 surface expression. Further correlation tests between these lipid species showed that they were all significantly and positively correlated with each other (Figure 5F).

Finally, I sought to identify critical resting platelet and releasate lipids that explained the reactivity of platelets, indicated by net increase in surface PAC1 and P-selectin expressions. To achieve this, Pearson correlation tests were performed between the resting platelet and releasate lipidomes and the net increment to PAC1 and P-selectin levels on platelet surface post collagen/thrombin stimulation. I identified resting platelet LPC-O as the only lipid class that significantly and positively correlated with collagen-induced increment to platelet surface PAC1 (Figure 5G). Similarly, only resting platelet PE 38:7 showed a very strong tendency (corrected p-value = 0.076) toward negative correlation with thrombin-induced net increase in surface P-selectin expression (Figure 5H).

## Discussion

Lipid has long been shown to play pivotal roles in human platelet activation and to be involved in platelet related diseases^11–15^. Similar to earlier platelet lipidomics studies, our results indicate the occurrence of the following events during platelet activation: (1) increased phospholipases, e.g., cytosolic phospholipase A (cPLA), phospholipase C activities, as indicated by phospholipid utilization (e.g., PI 38:4) as the substrate and the release of AA and other PUFAs, DGs and PAs. (2) enhanced production of lipid mediators utilizing the liberated PUFA species. (3) release of the prothrombotic lipids such as the lipid mediators, S1P, DG and a number of distinctive PLs^5–8,15–17^. Despite differences in study cohorts, analytical platforms and lab environments, some less mentioned outcomes reported earlier are likewise found in the current study. For instance, similar to the report from Peng, et al., I observed increase in some platelet lyso-PL classes only occurring to collagen but not thrombin stimulation (Figure 3E-H; discussed later)^7,18^. Likewise, reductions to several TG species in activated platelets with thrombin but not collagen treatment was observed (likely being consumed due to a higher energy demand), and the increment to several polyunsaturated PA species (indicative of AA re-esterification occurring specifically to this PL class) (Figure 3H) ^7^. These similarities not only corroborate earlier findings, but also indicate changes in platelet lipidomes in response to stimulations involve tightly regulated reactions which are highly conserved among human beings.

Our study mapped alterations to platelet and releasate lipidomes in response to thrombin and collagen treatments, in which I not only identified lipid classes that have not been reported in earlier releasate lipidomics studies, e.g., LPC(O), LPC(P), LPE(P), CL, but also presented the data in a comprehensive and intuitive manner (Figure 4 & Supplementary Table S4) and thus providing a unique angle to deciphering platelet-releasate relationship at the lipidome level^8,19^. One finding that to my knowledge has never been reported before is that acute collagen treatment, at least with the concentration tested here, resulted in the retention of a large number of lipid species while still activating cPLA2 (indicated by increased platelet lyso-PLs) and the production and/or release of lipid mediators (TXB2, 12-HETE and 8(9)-EET), LPI and S1P (Figure 3H & 4A). Given that no microparticles (granules, extracellular vesicles, etc.) were further isolated from releasate in this study, it is unclear if this was caused by impeded degranulation. This finding also partially explains the higher lyso-PL levels (with LPC-O and LPE being most apparent) in collagen-treated platelets than the thrombin treated ones, even though the latter more strongly activated phospholipases and led to more pronounced PL degradation (Figure 3)^7^. This observation was also supported by the low and close-to-baseline expression of platelet surface P-selectin, which has often been regarded as an indicator for platelet degranulation, similar to earlier studies (Figure 5A)^9,20^. Reserving instead of releasing the lipid contents may be one mechanism behind collagen not stimulating platelets as strongly as thrombin, since many of the prothrombotic lipid species were not released and therefore could not further instigate platelet activation. Thrombin treatment, especially with a prolonged incubation period, triggered changes in the lipid profile resembling platelets undergoing senescence, such as the increase in certain ceramide species and the release of PS^21^. Indeed, several studies have shown that thrombin not only acts as an inducer for platelet activation, but can also initiate apoptosis, which likely explains the observed alterations to platelet lipid profile here^20,22,23^. Intriguingly, many of the acutely produced and released lipids known to take crucial parts in thrombogenesis, such as LM, LPI and S1P, did not increase and/or be released further with prolonged treatment. By contrast, the levels of several apoptosis-related lipids (e.g., ceramide, PA, PE, PS, DG), which also have been shown to be thrombogenic, were generally secreted further at the late stage (Figure 3E-G)^8,24–31^. This result suggests that the effect of thrombin, at least for its impacts on thrombogenic lipid production and secretion, may be time dependent, with its ability to trigger platelet apoptosis and the release of a different set of (vs acute treatment) lipid species to further stimulate thrombogenesis occurring at the later stage. This study further echoes the pivotal roles apoptosis and the release of proapoptotic platelet contents can play in many thrombotic diseases^32–34^ and offers potential explanations from the lipid perspective.

Slatter et al. demonstrated distinctive basal and stimulated lipidomes and the different levels of platelet activation among genetically unrelated donors. However, no decisive conclusion was drawn due to the very small sample size in that study (n=3)^6^. Here I was able to exploit platelet surface expression of P-selectin (indicator for degranulation/chemotaxis) and PAC1 (targeting activated glycoprotein IIb/IIIa complex)^9^ to further interrogate the relationship between platelet lipidome and activation. Di- and polyunsaturated PC and PE were identified to be negatively correlated with PAC1 expression upon collagen stimulation (Figure 5F). While the negative correlation between PAC1 and PC 40:5 and PE 40:5 may be attributable to more active phospholipase activity, which utilized polyunsaturated (mostly arachidonyl) phospholipid as its substrate, this mechanism is less relevant to the di-unsaturated (36:2 and 38:2) PC and PE species. Earlier studies found that phospholipid is crucial to the stability and shape of platelet membrane and could impact platelet activation and degranulation^35^. Moreover, it has been shown that the platelets from severely obese individuals, which were likely hyperreactive and hypercoagulable^36^, had slightly but significantly less PC and PE^37^. Yet another study showed several short-chained, saturated and monounsaturated PC and PE species being strongly upregulated in acute coronary syndrome, with concomitant decrease in several long chain and polyunsaturated PC and PE species. Among these PL species, PC 8:0_10:0 was further identified as a strong thrombogenic agent^11^. Taken together, it appears that long-chained and unsaturated PLs have the tendency to stabilize platelet from the effect of prothrombogenic short-chained and saturated PL species, which could explain the negative correlations observed in our study. Other than PC and PE, CoQ was the only lipid class that negatively correlated with platelet surface PAC1. CoQ is a mitochondrial electron transport chain component and is also a potent antioxidant^38^. An earlier study showed that platelet CoQ level was lower among rheumatoid arthritis patients, which was linked to platelet mitochondrial dysfunction and increased ROS production^39^. Furthermore, in vitro and animal studies have shown that CoQ supplement suppressed both integrin αIIbβ3 inside-out and outside-in signaling, lowered ROS level and could hinder platelet activation in reaction to different stimuli^40,41^. Given that PAC1 is a monoclonal antibody that targets activated αIIbβ3, our data again suggest that CoQ may modulate platelet activation via targeting this specific integrin on platelet membrane (Figure 5F).

One of the most intriguing questions being interrogated in this study was whether I could predict the reactivity of platelets from one donor prior to stimulations, a query that bears clinical significance as there are benefits if diagnostic assays requiring live and freshly isolated platelets of relatively large quantities and an extra platelet activation step (e.g., aggregometry) can be substituted with lipid profiling that only requires small amounts of frozen platelet lysates and without the need for platelet activation. Although strictly speaking the Rest group (representing the unstimulated platelets) used in our study were incubated for an addition 10 min, these samples should accurately reflect the “pre-activation” state. To our knowledge, one study by Vadaq et al. utilized a similar approach, in which they correlated plasma metabolites to platelet reactivity, which was proposed to have gut microbiota origin and reflected the extracellular influence on platelets^42^. Our lipidomics study, on the other hand, elucidated the impact of platelet lipid on its own reactivity (intracellular) and identified baseline LPC-O and PE 38:7, respectively, as potential lipid predictors for collagen-stimulated PAC1 expression and thrombin-stimulated P-selectin expression (Figure 5G&H). LPC-O, also known as “lyso-platelet activating factor”, is a precursor for platelet activating factor (PAF, acetyl-glyceryl-ether-phosphorylcholine), a lipokine that strongly activates platelets by acting on surface GPCR PAF receptors, via the remodeling pathway. Having a higher level of intra-platelet LPC-O available may result in more rapid production and release of PAF upon stimulation, as PAF is not stored in resting cells. Furthermore, the remodeling pathway has generally been considered as the major pathway for PAF production in an inflammatory setting, which further supports this hypothesis. Alternatively, the differential resting platelet LPC-O levels among the donors may reflect cPLA2 activity, a crucial platelet lipolytic enzyme that not only catalyses the essential first step for PAF synthesis but also other reactions closely related to platelet activation, such as the liberation of AA from phospholipid^43–45^, and hence the strong correlation with PAC1 expression. Notably, our correlation analysis between platelet and releasate suggests that LPC-O could be secreted, at least in the case of thrombin stimulation (Figure 4G), similar to an earlier report^46^. Furthermore, studies have demonstrated that PAF receptor could be activated (to a lesser extent) by PAF-like phospholipid bearing certain structural similarities^43,47^, although for LPC-O this effect has been controversial^48,49^. Additionally, one study showed that exogenous LPC-O directly supplemented PAF production and rescued the inhibition of cPLA2 by adenosine and histamine^50^. Further correlation analysis nonetheless suggests that the secreted LPC-O should not play a major role here, as platelet and releasate LPC-O only correlate modestly (R^2^ = 0.29) and the latter also does not correlate with PAC1 expression induced by collagen at 10 min (R^2^ = 0.03). The relationship between resting platelet PE 38:7 and thrombin-stimulated P-selection expression is less clear. Studies suggest that disturbance to inner membrane aminophospholipid (PE and PS) occurs during platelet activation^15^. A number of studies also demonstrated that supplementing polyunsaturated FA, which was incorporated into membrane phospholipid, mitigated platelet activation^51,52^. Furthermore, a small DHA (22:6n3) and EPA supplement cohort study demonstrated human platelets were able to incorporate these two polyunsaturated FA, with DHA supplement significantly suppressed various markers for platelet activation, including platelet surface P-selectin expression^53^. Similar effects of DHA and EPA have also been shown to lower soluble P-selectin^54^. Having a higher level of PE 38:7 in cell and/or alpha granule membrane may prevent P-selectin from surfacing and subsequent degranulation. Given that ω-3, ω-6 or even individual PUFA species could exert dissimilar effects, further studies such as clarifying the fatty acyl composition of PE 38:7 will be required to decipher the underlying mechanisms behind this strong negative correlation between PE 38:7 and platelet P-selectin expression^52^.

In conclusion, our platelet lipidomics study provides a comprehensive view regarding alterations to platelet and releasate lipid profiles that occur during acute and chronic stimulations with collagen and thrombin. Detailed mappings of platelet and releasate lipidomes were carried out. One lipid class, LPC-O, and one lipid species, PE 38:7, have been identified to potentially predict the reactivity of platelet to stimulation to collagen and P-selectin, which may find application in predicting platelet hyperreactivity that may occur in thrombotic disorders.

## Methods

### Reagents & Materials

Collagen was purchased from Helena Laboratories (TX, USA); FITC-conjugated PAC1 and APC-conjugated anti-P-selectin (CD62p) were obtained from BD Biosciences (NSW, AU); LC/MS grade acetonitrile, isopropanol and methanol and HPLC glass vials and caps were from Thermo Fisher Scientific (MA, USA); the rest were acquired from Sigma Aldrich (NSW, AU).

### Washed human platelets

Human platelets were isolated from blood collected following the *National Statement on Ethical Conduct in Human Research (2007) – updated 2018* guideline and was approved by the Human Research Ethics Committees of the University of Sydney 2014/HE000244. Human blood was collected intravenously from 15 healthy donors in ACD tubes. Blood cell counts in whole blood were determined with an automatic haematology analyser (Sysmex). Washed platelets suitable for lipidomics analysis were prepared using the centrifugation method following protocols from earlier studies ^7,15,55^. Briefly, 30-50 mL whole blood was first spun 200g, room temperature, brake = 0 for 20 min to obtain platelet rich plasma (PRP). PRP was then added to warm (37°C) HEPES Tyrode buffer (pH 6.5) containing 5 mM glucose and 10 % ACD to 1:4 (v/v) ratio and left in a 37°C water bath for 30 min. Subsequently, another 800g, room temperature and brake = 0 spin for 5 min was applied to pellet platelets. The platelets were then resuspended in 300-600 μL warm (37°C) HEPES Tyrode buffer (pH 7.4) containing 5 mM glucose and left for 30-60 min in a 37°C water bath to allow dispersion^15^. Resuspended platelets then underwent cell number adjustment to make the final cell counts 4*10^5^/μL with the addition of warm HEPES Tyrode glucose buffer (pH 7.4).

### Ex vivo platelet activation

200 μL of the above washed platelets were first preconditioned with CaCl_2_ (1 mM) in a 37°C water bath for 10 min. Wash platelets were then either underwent further incubation uninterrupted (resting), or treated with 0.2 or 1.0 U/mL thrombin or 10 μg/mL collagen for 10 or 60 min in a 37°C water bath. Afterwards, platelets and releasates were separated by 770g, 5 min, soft brake centrifugation in a 37°C warm room. Supernatants (releasates) from the samples were then swiftly collected and the platelets and releasates were quickly stored in a −80°C freezer.

### Flow Cytometry

10^6^ platelets were suspended in 100 µL phosphate buffer saline (PBS) under room temperature, followed by the addition of 2 µL anti-PAC1-FITC (BD, NSW, AU) or anti-CD62p-APC (BD, NSW, AU). An unstained sample was also prepared to control the background signal. Straight after the addition of antibodies, platelets were then not stimulated (resting) or stimulated with 0.2 U/mL thrombin or 10 μg/mL collagen for 10 min before the reaction were terminated by adding 300 µL of HEPES Tyrode buffer. Platelets were then analysed with a BD FACS Accuri flow cytometer. 10^4^ gated events were collected with channels corresponding to the conjugated fluorophores (FL1-A for anti-PAC1-FITC; FL4-A for anti-CD62p-APC).

### Platelet and releasate lipid extractions

A 1-butanol/methanol (BUME) single phase lipid extraction method was used for platelet and releasate lipid extraction^10^. Prior to sample preparation, the sample order has been randomized using Excel (Microsoft). For platelet samples, 10^6^ platelets were lysed in 50 μL Tris-HCl with 100 µM BHT by probe sonication. BCA assays were subsequently performed to determine platelet lysates protein concentrations, which were then adjusted to 1 g/L. 10 μL of the platelet lysates were taken and mixed with 15 μL of internal standard mix (Supplementary Table S5) and 35 μL BUME solvent (1-butanol/methanol = 1:1 with 5 mM ammonium formate), followed by vortex and water bath sonication under room temperature for 1 hr. Afterwards, the samples underwent 16100g, room temperature centrifugation for 10 min and the supernatants were transferred to 1.5 mL glass vials with fused inserts for lipidomic analysis. For releasate, 100 μL was taken from each sample and mixed with 15 μL of internal standard mix and 485 μL BUME solvent, followed by vortex and water bath sonication under room temperature for 1 hr. Afterwards, the samples were centrifuged 16100g for 10 min (room temperature). Supernatants were collected and dried in a SpeedVac system (Thermo Fisher Scientific, MA, USA). Dried samples containing extracted lipid were reconstituted in mixture of 50 μL BUME solvent and 10 μL H_2_O and then water bath sonicated for 10 min. Samples were then centrifuged 16100g, room temperature, before the supernatants were collected and transferred to 1.5 mL glass vials with fused inserts for lipidomic analysis.

### Targeted lipidomics

Platelet and releasate lipid profiles were determined using a purposely built targeted LC/MS-MS method on a TSQ Altis triple quadrupole mass spectrometer equipped with a H-ESI source and coupled with a Vanquish Horizon liquid chromatography (LC) system (Thermo Fisher Scientific, MA, USA). Briefly, autosampler injection volume was 5 μL and temperature was set 10°C. Liquid chromatographic separation was achieved with a 100*2.1 mm Supelco Ascentis® Express C18, 2 μm UHPLC Column (Sigma Aldrich, NSW, AU) attached to a SecurityGuard™ ULTRA C18 guard column (Phenomenex, CA, USA), with 35°C column oven temperature. A binary gradient buffer system was used as the mobile phase throughout the 23 min run time. Solvent A comprised acetonitrile/H_2_O = 6:4, 10 mM ammonium formate, 5 μM phosphoric acid and 0.1% (v/v) formic acid; solvent B comprised isopropanol/acetonitrile/H_2_O = 90:9:1, 10 mM ammonium formate and 0.1% (v/v) formic acid. The set gradient was: 40% B, 0 min −> 100% B, 20-21.5 min −> 40% B, 22-23 min. The flow rate was mostly fixed at 0.3 mL/min, except for one section of the re-equilibration stage (21.5-22 min) during which it was increased to 0.38 mL/min and subsequently returned to 0.3 mL/min at 22.8 min.

Positive and negative scheduled single reaction monitoring (SRM) modes were applied to acquire targeted lipidomic data. SRM transition tables were generated with the following tools: triacylglycerol SRMs were created and identified using TAILOR-MS^56^ (Supplementary Table S1); Positive cardiolipin, acylcarnitine and coenzyme Q SRM transitions were obtained/modified from earlier publications^57,58^; transitions for the remaining lipid classes were obtained from LipidCreator database^59^. Pooled platelet and releasate samples were used for several pilot runs to verify the existence of the *in silico* generated lipid SRMs before platelet and releasate batch runs. To distinguish ether and ester phospholipid and lysophospholipid peaks and to exclude isotopic sphingomyelin peaks, separate human platelet lipid analysis was carried out on a high mass resolution Q Exactive HFX Orbitrap instrument coupled with Vanquish Horizon LC (Thermo Fisher Scientific, MA USA) using identical LC settings to generate matched retention times to the targeted analysis. Plasmanyl/plasmenyl PC and PE peak determination was achieved with acid treatments^60^. 19 deuterated or non-physiological lipid standards were used to optimize collision energies and RF-Lens settings at lipid class level, and as spike-internal standards (refer to the “Lipid extraction” section) for semi-quantification. Among them, 14 were constituents of SPLASH® LIPIDOMIX® Mass Spec Standard (Avanti Research, AL, USA) and the additional 5 were Cer(d18:1-d7/15:0), Glucosyl(β) Cer(d18:1/17:0), FA(15:0), CL(14:0/14:0/14:0/14:0) and Sph(d17:1) (Sigma Aldrich, NSW, AU). Totally, transitions representing 549 lipid species encompassing 32 lipid classes were detected in platelets and those representing 562 lipid species encompassing 34 lipid classes were identified in releasates (Supplementary Table S5 & S6). Calculated coefficient of variation distributions for QC samples, extraction efficiencies and matrix effects can be found in Supplementary Table S7-S9.

### Data Analysis

FACS data were analysed using the Python package Cytoflow^61^. Raw LC/MS-MS data examination, deconvolution and manually peak integration (when necessary) were performed using Skyline^62^. The exported spreadsheets containing peak areas were subsequently tidied and converted to concentrations using in-house Python scripts utilizing Numpy and Pandas packages^63,64^. Statistical analysis was carried out using Python Scipy, Statsmodels and Pingouin packages^65–67^. For effects of different stimulations and treatment time on platelet and releasate, two-way repeated measure ANOVA was first performed on log10-transformed concentration data (log nmol/mg P for platelet; log noml/mL for releasate; Greenhouse-Geisser corrected p-values), followed by paired t-tests as *post hoc* analysis. Pearson correlation tests were used to analyse “lipid vs lipid” and “lipid vs platelet surface marker” relationships. All p-values were further corrected for multiple comparisons with Benjamini-Hochberg approach. For flow cytometry data, Mann-Whitney-Wilcoxon tests followed by Bonferroni corrections were performed. P-values < 0.05 were considered statistically significant, while p-values 0.05 - 0.06 were deemed having a strong tendency toward significance. Python Matplotlib, Seaborn and NetworkX packages were used for plot generation^68–70^.

## Acknowledgements

The author would like to express his greatest gratitude toward Associate Professor Freda Passam for providing samples, equipment, consumables and kindly allowing him to complete this study outside of his normal work duty. The author wishes to also thank Professor Anthony Don, Dr Ben Crossett and Dr Lake-Ee Quek for their valuable comments. Furthermore, the author thanks all the blood donors who participated in this study by donating blood. The author declares no conflict of interest.

